# Evolutionary transcriptomics implicates *HAND2* in the origins of implantation and regulation of gestation length

**DOI:** 10.1101/2020.06.15.152868

**Authors:** Mirna Marinić, Katelyn Mika, Sravanthi Chigurupati, Vincent J. Lynch

## Abstract

The developmental origins and evolutionary histories of cell types, tissues and organ systems contribute to the ways in which their dysfunction leads to disease. In mammals for example, the nature and extent of maternal-fetal interactions, how those interactions develop, and their evolutionary history likely influence diseases of pregnancy such as infertility and preterm birth. Here we show genes that evolved to be expressed at the maternal-fetal interface in Eutherian (‘Placental’) mammals play essential roles in the evolution of pregnancy and are associated with immune system disorders and preterm birth. Among these genes is the transcription factor *HAND2*, which suppresses estrogen signaling, an innovation of Eutherians, thereby allowing blastocyst implantation. We found that *HAND2* is dynamically expressed in the decidua throughout the menstrual cycle and pregnancy, gradually decreasing to reach a low at term. HAND2 regulates a small but distinct set of target genes in endometrial stromal fibroblasts including the cytokine *IL15*, which was also dynamically expressed throughout the menstrual cycle and gestation, and promoted the migration of natural killer cells and extravillous cytotrophoblasts. Remarkably, we found that the *HAND2* promoter loops to a distal enhancer containing SNPs implicated in the regulation of gestation length and birth weight. Collectively, these data connect *HAND2* expression at the maternal-fetal interface with the evolution of implantation and gestation length regulation, and preterm birth.

## Introduction

The ontogeny and evolutionary history of cell types, tissues and organ systems, as well as the life histories of organisms biases the ways in which dysfunctions in those systems underlie disease (Varki, 2012). Thus a mechanistic understanding of how cells, tissues and organs evolved their functions, and how organism’s life histories influence them, may provide clues to the molecular etiologies of disease. The most common way of utilizing evolutionary information to characterize the genetic architecture of disease is to link genetic variation within a species to phenotypes using quantitative trait loci (QTL) or genome wide association studies (GWAS). An alternative approach is to identify fixed genetic differences between species that are phylogenetically correlated with different disease relevant phenotypes. While the risk of cancer increases with age across mammals, for example, the prevalence of cancer types varies by species (Abegglen *et al*., 2015), likely because of differences in genetic susceptibility to specific cancers, structure of organ and tissue systems, and life exposures to carcinogens (Varki and Varki, 2015). Similarly, the risk of cardiovascular disease (CVD) increases with age across species, but the pathophysiology of CVD can differ even between closely related taxa such as humans, in which CVD predominantly results from coronary artery atherosclerosis, and the other Great Apes (Hominids), in which CVD is most often associated with interstitial myocardial fibrosis (Varki *et al*., 2009).

Extant mammals span major stages in the origins and diversification of pregnancy, thus a mechanistic understanding of how pregnancy originated and diverged may provide unique insights into the ontogenetic origins of pregnancy disorders. The platypus and echidna (Monotremes) are oviparous, but the embryo is retained in the uterus for 10–22 days, during which the developing fetus is nourished by maternal secretions delivered through a simple placenta, prior to the laying of a thin, poorly mineralized egg that hatches in ∼2 weeks (Hill, 1936). Live birth (viviparity) evolved in the stem-lineage of Therian mammals, but Marsupials and Eutherian (‘Placental’) mammals have dramatically different reproductive strategies. In Marsupials, pregnancies are generally short (∼25 days) and completed within the span of a single estrous cycle (Renfree and Shaw, 2001; Renfree, 2010). Eutherians, in contrast, evolved a suite of traits that support prolonged pregnancies (up to 670 days in African elephant), including an interrupted estrous cycle, which allows for gestation lengths longer than a single reproductive cycle, maternal-fetal communication, maternal recognition of pregnancy, implantation of the blastocyst and placenta into uterine tissue, differentiation (decidualization) of endometrial stromal fibroblasts (ESFs) in the uterine lining into decidual stromal cells (DSCs), and maternal immunotolerance of the antigenically distinct fetus, i.e. the fetal allograft (Guleria and Pollard, 2000; Moffett and Loke, 2004; Erlebacher, 2013).

Gene expression changes at the maternal-fetal interface underlie evolutionary differences in pregnancy (Hou *et al*., 2012; Lynch *et al*., 2015; Armstrong *et al*., 2017), and thus likely also pathologies of pregnancy such as infertility, recurrent spontaneous abortion (Kosova *et al*., 2015), preeclampsia (Elliot, 2017; Arthur *et al*., 2018; Varas Enriquez, McKerracher and Elliot, 2018) and preterm birth (Plunkett *et al*., 2011; Swaggart, Pavlicev and Muglia, 2015; LaBella *et al*., 2019). Here, we assembled a collection of gene expression data from the pregnant/gravid maternal-fetal interface of tetrapods and used evolutionary methods to reconstruct gene expression changes during the origins of mammalian pregnancy. We found that genes that evolved to be expressed at the maternal-fetal interface in the Eutherian stem-lineage were enriched for immune functions and diseases, as well as preterm birth. Among the recruited genes was the transcription factor *Heart-and neural crest derivatives-expressed protein 2* (*HAND2*), which plays essential roles in neural crest development (Srivastava *et al*., 1997), cardiac morphogenesis (Srivastava *et al*., 1997; Shen *et al*., 2010; Tamura *et al*., 2013; Lu *et al*., 2016; Sun *et al*., 2016), and suppressing estrogen signaling during the period of uterine receptivity to implantation (Huyen and Bany, 2011; Li *et al*., 2011; Shindoh *et al*., 2014; Fukuda *et al*., 2015; Mestre-Citrinovitz *et al*., 2015; Murata *et al*., 2019; Šućurović *et al*., 2020). We determined that *HAND2* expression at the first trimester maternal-fetal interface was almost entirely restricted to cell types in ESF lineage, and is regulated by multiple transcription factors that control progesterone responsiveness. Moreover, the *HAND2* promoter loops to an enhancer with single nucleotide polymorphisms (SNPs) that have been implicated by GWAS in the regulation of gestation length (Warrington *et al*., 2019; Sakabe *et al*., 2020). Furthermore, we showed that HAND2 regulates interleukin 15 (*IL15*) expression in ESFs, and that ESF-derived IL15 influences the migration of natural killer and trophoblast cells. These data suggest that HAND2 and IL15 signaling played an important role in the evolution of implantation and regulation of gestation length.

## Results

### Genes that evolved endometrial expression in Eutherian mammals are enriched in immune functions

We previously used comparative transcriptomics to reconstruct the evolution of gene expression at the maternal-fetal interface during the origins of mammalian pregnancy (Lynch *et al*., 2015). Here, we assembled a collection of new and existing transcriptomes from the pregnant/gravid endometria of 15 Eutherian mammals, three Marsupials, one Monotreme (platypus), two birds, six lizards, and one amphibian (**Supplementary Table 1**). The complete dataset includes expression information for 21,750 genes from 28 species. Next, we transformed continuous transcript abundance estimates values into discrete character states such that genes with Transcripts Per Million (TPM) ≥ 2.0 were coded as expressed (state=1), genes with TPM < 2.0 were coded as not expressed (state=0), and genes without data in specific species were coded as missing (state=?). We then used parsimony to reconstruct ancestral transcriptomes and trace the evolution of gene expression gains (0 → 1) and losses (1 → 0) in the endometrium.

**Table 1.**
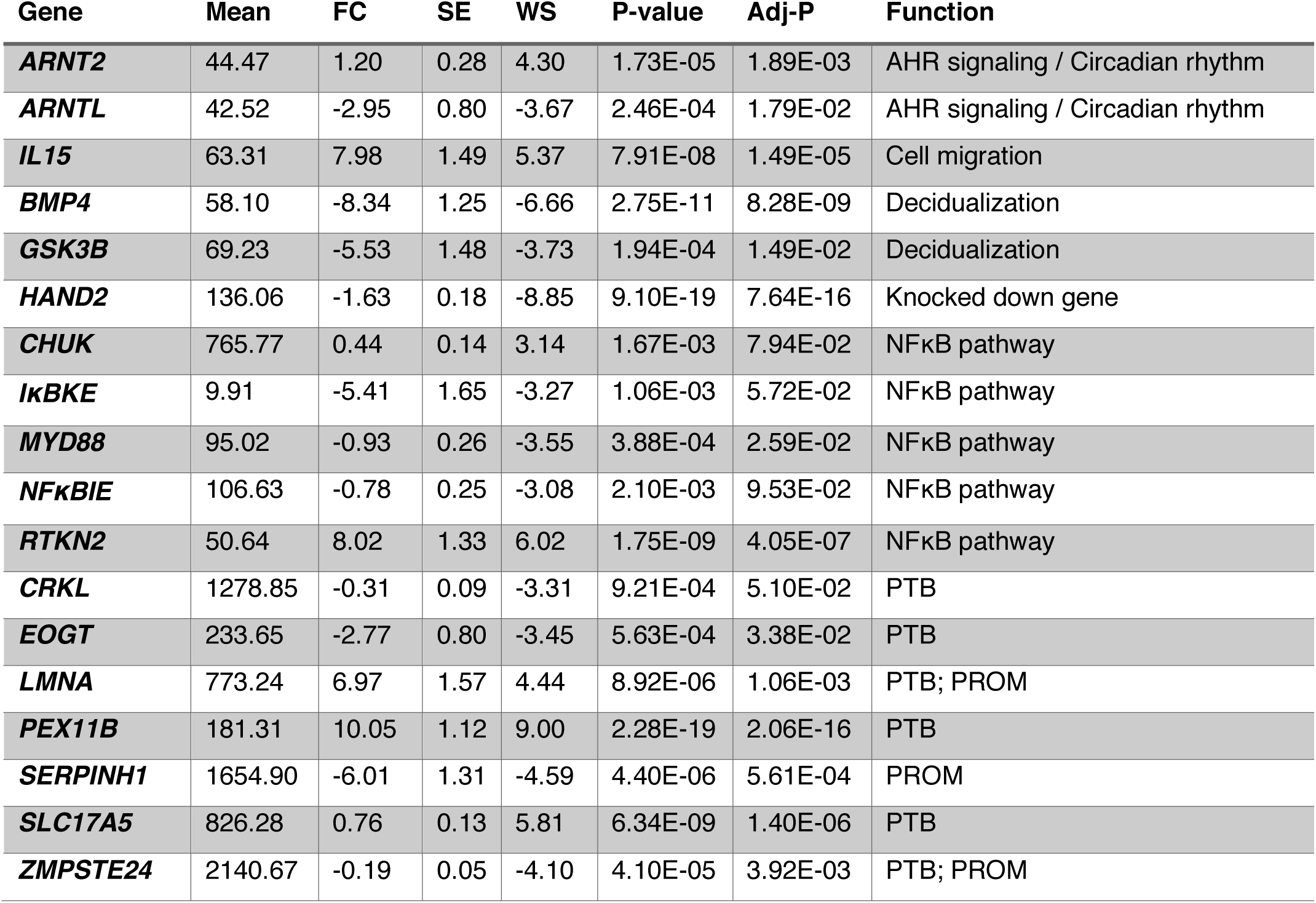
Exemplar genes differentially expressed in ESFs upon siRNA-mediated *HAND2* knockdown. Mean, base mean expression level. FC, log_2_ fold change. SE, standard error in log_2_ fold change. WS, Wald statistic. P-value, Wald test P-value. Adj-P, Benjamini-Hochberg (BH) adjusted P-value. Function, function of gene inferred from WikiPathway 2019 human annotation. Association with preterm birth (PTB, HP:0001622) and premature rupture of membranes (PROM, HP:0001788) inferred from human phenotype ontology annotation.

| Gene | Mean | FC | SE | WS | P-value | Adj-P | Function |
| --- | --- | --- | --- | --- | --- | --- | --- |
| ***ARNT2*** | 44.47 | 1.20 | 0.28 | 4.30 | 1.73E-05 | 1.89E-03 | AHR signaling / Circadian rhythm |
| ***ARNTL*** | 42.52 | -2.95 | 0.80 | -3.67 | 2.46E-04 | 1.79E-02 | AHR signaling / Circadian rhythm |
| ***IL15*** | 63.31 | 7.98 | 1.49 | 5.37 | 7.91E-08 | 1.49E-05 | Cell migration |
| ***BMP4*** | 58.10 | -8.34 | 1.25 | -6.66 | 2.75E-11 | 8.28E-09 | Decidualization |
| ***GSK3B*** | 69.23 | -5.53 | 1.48 | -3.73 | 1.94E-04 | 1.49E-02 | Decidualization |
| ***HAND2*** | 136.06 | -1.63 | 0.18 | -8.85 | 9.10E-19 | 7.64E-16 | Knocked down gene |
| ***CHUK*** | 765.77 | 0.44 | 0.14 | 3.14 | 1.67E-03 | 7.94E-02 | NFκB pathway |
| ***IκBKE*** | 9.91 | -5.41 | 1.65 | -3.27 | 1.06E-03 | 5.72E-02 | NFκB pathway |
| ***MYD88*** | 95.02 | -0.93 | 0.26 | -3.55 | 3.88E-04 | 2.59E-02 | NFκB pathway |
| ***NFκBIE*** | 106.63 | -0.78 | 0.25 | -3.08 | 2.10E-03 | 9.53E-02 | NFκB pathway |
| ***RTKN2*** | 50.64 | 8.02 | 1.33 | 6.02 | 1.75E-09 | 4.05E-07 | NFκB pathway |
| ***CRKL*** | 1278.85 | -0.31 | 0.09 | -3.31 | 9.21E-04 | 5.10E-02 | PTB |
| ***EOGT*** | 233.65 | -2.77 | 0.80 | -3.45 | 5.63E-04 | 3.38E-02 | PTB |
| ***LMNA*** | 773.24 | 6.97 | 1.57 | 4.44 | 8.92E-06 | 1.06E-03 | PTB; PROM |
| ***PEX11B*** | 181.31 | 10.05 | 1.12 | 9.00 | 2.28E-19 | 2.06E-16 | PTB |
| ***SERPINH1*** | 1654.90 | -6.01 | 1.31 | -4.59 | 4.40E-06 | 5.61E-04 | PROM |
| ***SLC17A5*** | 826.28 | 0.76 | 0.13 | 5.81 | 6.34E-09 | 1.40E-06 | PTB |
| ***ZMPSTE24*** | 2140.67 | -0.19 | 0.05 | -4.10 | 4.10E-05 | 3.92E-03 | PTB; PROM |

We identified 958 genes that potentially evolved endometrial expression in the Eutherian stem-lineage (**Figure 1A**), including 149 that unambiguously evolved endometrial expression (**Supplementary Table 2**). These 149 genes were significantly enriched in pathways related to the immune system (**Supplementary Table 3**), although only two pathways were enriched at False Discovery Rate (FDR) ≤ 0.10, namely, “Cytokine Signaling in Immune System” (hypergeometric *P*=1.97×10^−5^, FDR=0.054) and “Signaling by Interleukins” (hypergeometric *P*=4.09×10^−5^, FDR=0.067). Unambiguously recruited genes were also enriched in numerous human phenotype ontology terms (**Supplementary Table 4**) but only two, “Immune System Diseases” (hypergeometric *P*=3.15×10^−8^, FDR=2.52×10^−4^) and “Preterm Birth” (hypergeometric *P*=4.04×10^−4^, FDR=8.07×10^−4^), at FDR ≤ 0.10. In contrast, these genes were enriched in numerous biological process gene ontology (GO) terms (**Supplementary Table 5**), nearly all of which were related to regulation of immune system, including “Leukocyte Migration” (hypergeometric *P*=1.17×10^−7^, FDR=1.29×10^−3^), “Inflammatory Response” (hypergeometric *P*=8.07×10^−7^, FDR=2.21×10^−3^), and “Cytokine-mediated Signaling Pathway” (hypergeometric *P*=2.18×10^−5^, FDR=0.013).

**Figure 1.**
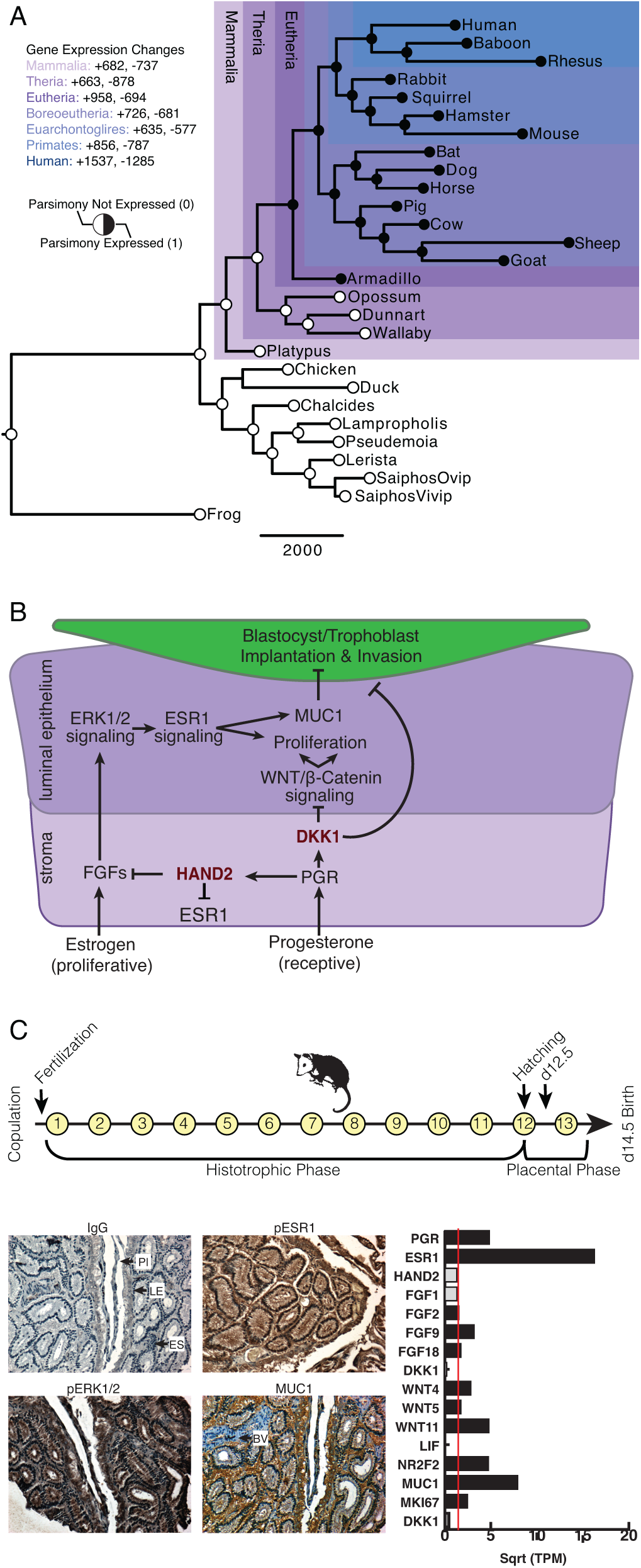
Recruitment of HAND2-mediated anti-estrogenic signaling in the Eutherian endometrium. A. Evolution of *HAND2* expression at the maternal-fetal interface. Amniotes phylogeny with horizontal branch lengths drawn proportional to the number of gene expression changes inferred by parsimony (most parsimonious reconstruction). Circles indicate *HAND2* expression in extant species and ancestral reconstructions. Black, expressed (state=1). White, not expressed (state=0). Inset legend shows the number of most gene expression changes from the root node to human (+ = gene expression gained; – = gene expression lost). B. Cartoon model of estrogen signaling and HAND2-mediated anti-estrogenic signaling in the endometrium. The estrogen-mediated signaling network is suppressed by progesterone through the activation of HAND2 and antagonists of canonical WNT/β-catenin mediated signaling pathways such as DKK1. In the proliferative phase of the reproductive cycle, estrogen acts through ESR1 in stromal cells to increase the production of fibroblast growth factors (FGFs), which serve as paracrine signals leading to sustained proliferation of epithelial cells. Active estrogen signaling maintains epithelial expression of Mucin 1 (MUC1), a cell surface glycoprotein that acts as a barrier to implantation. During the receptive phase of the cycle, however, progesterone induces HAND2 and DKK1 expression in the endometrial stroma, inhibiting production of fibroblast growth factors (FGFs), suppressing epithelial proliferation and antagonizing estrogen-mediated expression of MUC1, thereby promoting uterine receptivity to implantation. C. Upper, Timeline of pregnancy in *Monodelphis domestica* highlighting the histotrophic and placental phases. Bottom left, Immunohistochemistry showing phosphorylated ESR1 (pESR1), phosphorylated MAPK1/2 (pERK1/2), and MUC1 expression in paraffin embedded 12.5 d.p.c. pregnant opossum endometrium compared to control (IgG). Pl, placenta. LE, luminal epithelium. ES, endometrial stroma. BV, blood vessels. Bottom right, expression of genes related to HAND2-mediated anti-estrogenic signaling in RNA-Seq from 12.5 d.p.c. pregnant opossum endometrium. Data shown as square root (Sqrt) transformed Transcripts Per Million (TPM). The TPM=2 expression cutoff is shown as a red line; genes with TPM≥2 are shown as black bars.

### Recruitment of *HAND2* and anti-estrogenic signaling in Eutherians

Among the genes that unambiguously evolved endometrial expression in the Eutherian stem-lineage was the basic helix-loop-helix family transcription factor *Heart- and neural crest derivatives-expressed protein 2* (*HAND2*). *HAND2* plays an essential role in mediating the anti-estrogenic action of progesterone and the establishment of uterine receptivity to implantation (**Figure 1B**) (Huyen and Bany, 2011; Li *et al*., 2011; Fukuda *et al*., 2015; Mestre-Citrinovitz *et al*., 2015), suggesting an active role of this hormone during pregnancy in non-Eutherian mammals. To test this, we performed immunohistochemistry (IHC) on endometrial sections from day 12.5 of pregnancy (after the transition from the histotrophic to placental phase) in the short-tailed opossum (*Monodelphis domestica*). We observed strong staining for estrogen receptor alpha (*ESR1*; ERα) phosphorylated at serine 118 (a mark of transcriptionally active ERα) (Kato *et al*., 1995), phosphorylated ERK1/2, and MUC1 (**Figure 1C**). We also found that *ESR1, FGF2, FGF9, FGF18, MUC1*, several *WNT* genes that stimulate proliferation of the luminal epithelia, and *MKI67* (Ki-67), a mark of actively proliferating cells (Scholzen and Gerdes, 2000), were abundantly expressed in RNA-Seq data from day 12.5 pregnant *M. domestica* endometrium (**Figure 1C**). These data are indicative of persistent estrogen signaling during pregnancy in opossum.

### *HAND2* is expressed in endometrial stromal fibroblast lineage cells

To determine which cell types at the human maternal-fetal interface express *HAND2*, we used previously generated single-cell RNA-Seq (scRNA-Seq) data from first trimester decidua (Vento-Tormo *et al*., 2018). *HAND2* expression was almost entirely restricted to cell populations in the endometrial stromal fibroblast lineage (ESF1, ESF2, and DSC), with particularly high expression in ESF2s and DSCs (**Figure 2A**). HAND2 protein was localized to nuclei in ESF lineage cells in human pregnant decidua (**Figure 2B**) from Human Protein Atlas IHC data (Uhlén *et al*., 2015). We also used existing functional genomics data to explore the regulatory status of the *HAND2* locus (see Methods). Consistent with active expression, the *HAND2* locus in human DSCs is marked by histone modifications that typify enhancers (H3K27ac) and promoters (H3K4me3), and is located in a region of open chromatin as assessed by ATAC-, DNaseI- and FAIRE-Seq. Additionally, it is bound by transcription factors that establish endometrial stromal cell-type identity and mediate decidualization, including the progesterone receptor (PR), NR2F2 (COUP-TFII), GATA2, FOSL2, FOXO1, as well as polymerase II (**Figure 2C**). The *HAND2* promoter loops to several distal enhancers, as assessed by H3K27ac HiChIP data generated from a normal hTERT-immortalized endometrial cell line (E6E7hTERT), including a region bound by PR, NR2F2, GATA2, FOSL2 and FOXO1, that also contains SNPs associated with gestation length in recent GWAS (Warrington *et al*., 2019; Sakabe *et al*., 2020) (**Figure 2C**). *HAND2* was significantly up-regulated by decidualization of human ESFs into DSCs by cAMP/progesterone treatment (Log_2_FC=1.28, P=2.62×10^−26^, FDR=1.16×10^−24^), and significantly down-regulated by siRNAs targeting PR (Log_2_FC=−0.90, P=7.05×10^−15^, FDR=2.03×10^−13^) and GATA2 (Log_2_FC=−2.73, P=0.01, FDR=0.19) (**Figure 2D**). In contrast, siRNA-mediated knockdown of neither NR2F2 (Log_2_FC=−0.91, P=0.05, FDR=1.0) nor FOXO1 (Log_2_FC=0.08, P=0.49, FDR=0.74) significantly altered *HAND2* expression (**Figure 2D**).

**Figure 2.**
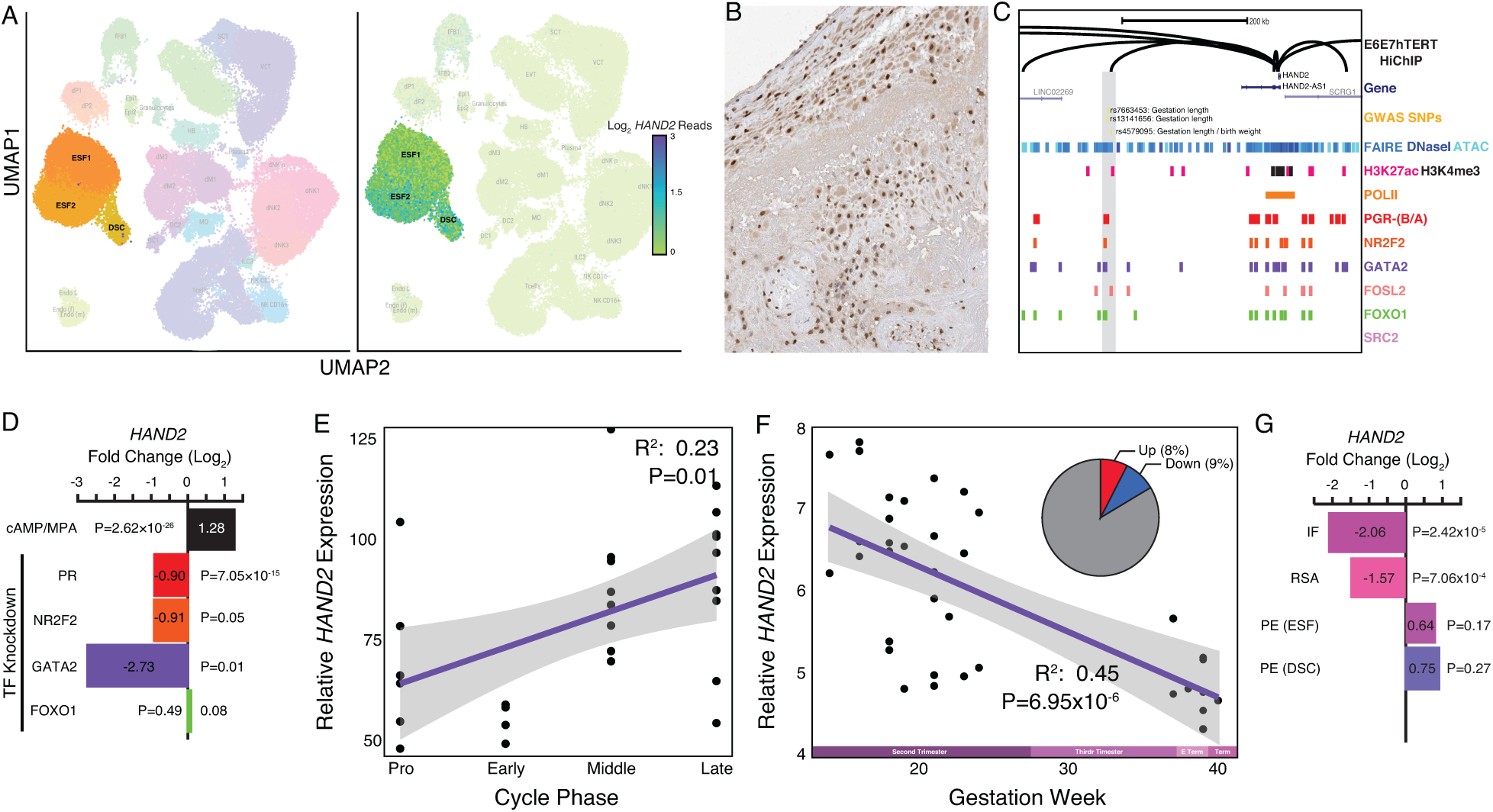
Expression of *HAND2* at the maternal-fetal interface. A. UMAP clustering of scRNA-Seq data from human first trimester maternal-fetal interface. Left, clusters colored according to inferred cell type. The ESF1, ESF2, and DSC clusters are highlighted. Right, cells within clusters are colored according to *HAND2* expression level. B. HAND2 protein expression in human pregnant decidua, with strong staining and localization in the nuclei of endometrial stromal cells (image color auto adjusted). Image credit: Human Protein Atlas. C. Regulatory landscape of the *HAND2* locus. Chromatin loops inferred from H3K27ac HiChIP, regions of open chromatin inferred from FAIRE-, DNaseI, and ATAC-Seq, and the locations of histone modifications and transcription factor ChIP-Seq peaks are shown. The location of SNPs associated with gestation length is also shown (highlighted in gray). Note that the *HAND2* promoter forms a long-range loop to a region marked by H3K27ac and bound by PR, NR2F2 (COUP-TFII), GATA2, FOSL2, and FOXO1. D. *HAND2* expression is up-regulated by in vitro decidualization of ESFs into DSC by cAMP/progesterone treatment, and down-regulated by siRNA-mediated knockdown of PR and GATA2, but not NR2F2 or FOXO1. n=3 per transcription factor knockdown. E. Relative expression of *HAND2* in the proliferative (n=6), early (n=4), middle (n=9), and late (n=9) secretory phases of the menstrual cycle. Note that outliers are excluded from the figure but not the regression; 95% CI is shown in gray. F. Relative expression of *HAND2* in the basal plate from mid-gestation to term (14-40 weeks); 95% CI is shown in gray. Inset, percent of up-(Up) and down-regulated (Down) genes between weeks 14-19 and 37-40 of pregnancy (FDR≤0.10). G. *HAND2* expression is significantly down-regulated in the endometria of women with implantation failure (IF, n=5) and recurrent spontaneous abortion (RSA, n=5) compared to fertile controls (n=5), but is not differentially expressed in ESFs or DSCs from women with preeclampsia (PE, n=5) compared to healthy controls (n=5).

### Differential *HAND2* expression throughout the menstrual cycle and pregnancy

Our observation that *HAND2* is progesterone responsive suggests it may be differentially expressed throughout the menstrual cycle and pregnancy. To explore this possibility, we utilized previously published gene expression datasets generated from the endometrium across the menstrual cycle (Talbi *et al*., 2006) and from the basal plate from mid-gestation to term (Winn *et al*., 2007). *HAND2* expression tended to increase from proliferative through the early and middle secretory phases, reaching a peak in the late secretory phase of the menstrual cycle (**Figure 2E**). In stark contrast, *HAND2* expression decreases throughout pregnancy reaching a low at term (**Figure 2F**). For comparison, 9% of genes were down-regulated between weeks 14-19 and 37-40 of pregnancy (FDR≤0.10). We also used previously published gene expression datasets (see Methods) to explore if *HAND2* was associated with disorders of pregnancy and found significant *HAND2* dysregulation in the endometria of women with infertility (IF) and recurrent spontaneous abortion (RSA) compared to fertile controls (**Figure 2G**). *HAND2* was not differentially expressed in ESFs or DSCs from women with preeclampsia (PE) compared to controls (**Figure 2G**).

### HAND2 regulates a distinct set of target genes

*HAND2* expression has previously been shown to play a role in orchestrating the transcriptional response to progesterone during decidualization in human and mouse DSCs (Huyen and Bany, 2011; Li *et al*., 2011; McConaha *et al*., 2011; Shindoh *et al*., 2014). However, whether *HAND2* has functions in other endometrial stromal lineage cells such as ESFs is unknown. Therefore, we used siRNA to knockdown *HAND2* expression in human hTERT-immortalized ESFs (T-HESC) and assayed global gene expression changes by RNA-Seq 48 hours after knockdown. We found that *HAND2* was knocked-down ∼78% (P=7.79×10^−3^) by siRNA treatment (**Figure 3A**), which dysregulated the expression of 553 transcripts (489 genes) at FDR ≤ 0.10 (**Figure 3A** and **Supplementary Table 6**). Genes dysregulated by *HAND2* knockdown had very little overlap with genes dysregulated by *PR, NR2F2, GATA2*, or *FOXO1* knockdown (**Figure 3B**). These data indicate that *HAND2* regulates a distinct set of target genes compared to transcription factors that establish cell-type identity and mediate the decidualization response. Indeed, only 30 genes are co-regulated by cAMP/progesterone, PR, GATA2, and HAND2 (**Figure 3B**).

**Figure 3.**
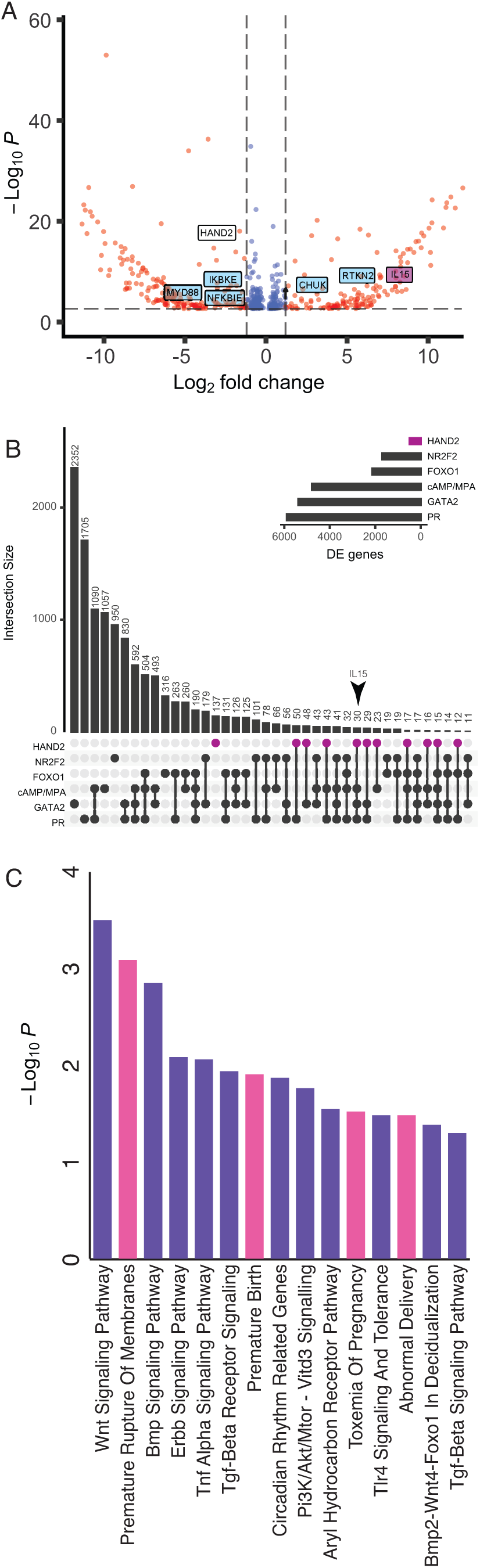
HAND2 regulates a distinct set of target genes, including *IL15*. A. Volcano plot of gene expression upon *HAND2* knockdown. Only genes that are significantly differentially expressed (DE) with FDR≤0.10 are shown. Genes with ≥1.5-fold changes in expression are shown in red, others in blue. B. UpSet plot showing intersection of genes DE by *HAND2, NR2F2, FOXO1, GATA2*, and *PR* knockdown (KD), and cAMP/MPA treatment. Inset plot shows number of DE genes for each treatment. Note large intersections for genes DE by cAMP/MPA/PR KD, GATA2 KD/PR KD, and cAMP/GATA2 KD/PR KD, but few intersections with HAND2 KD and other TF KDs. C. Pathways (purple) and human phenotype ontologies (pink) in which genes dysregulated upon *HAND2* KD are enriched.

Genes dysregulated by *HAND2* knockdown were enriched in several pathways and human phenotype ontologies relevant to endometrial stromal cells and pregnancy in general (**Figure 3C, Supplementary Table 7**, and **Supplementary Table 8**). Enriched pathways play a role in decidualization, such as “Wnt Signaling” (Peng *et al*., 2008; Hayashi *et al*., 2009; Sonderegger, Pollheimer and Knöfler, 2010; Franco *et al*., 2011; Wang *et al*., 2013), “BMP Signaling” (Ying and Zhao, 2000; Lee *et al*., 2007; Li *et al*., 2007; Wetendorf and DeMayo, 2012), “ErbB Signaling” (Lim, Dey and Das, 1997; Klonisch *et al*., 2001; Large *et al*., 2014), “TGF-beta Receptor Signaling” (Jones *et al*., 2006; Li, 2014; Ni and Li, 2017) and “BMP2-WNT4-FOXO1 Pathway in Human Primary Endometrial Stromal Cell Differentiation” (Gellersen and Brosens, 2003; Buzzio *et al*., 2006; Lee *et al*., 2007; Li *et al*., 2007; Brayer, Lynch and Wagner, 2009; Lynch *et al*., 2009; Kajihara, Brosens and Ishihara, 2013). Other enriched pathways included those involved in placental bed development disorders and preeclampsia (Chekir *et al*., 2006; Oliver *et al*., 2011; Guedes-Martins *et al*., 2013), the induction of pro-inflammatory factors via nuclear factor-κB (NFκB) through the “AGE/RAGE pathway” (Lappas, Permezel and Rice, 2007), the “Aryl Hydrocarbon Receptor Pathway”, which mediates maternal immunotolerance of the fetal allograft (Munn *et al*., 1998; Abbott *et al*., 1999; Funeshima *et al*., 2005; Hao *et al*., 2013), “Circadian Rhythm Related Genes”, which have been associated with implantation (Greenhill, 2014) and parturition (Roizen *et al*., 2007; Olcese, 2012; Olcese, Lozier and Paradise, 2013; Menon *et al*., 2016), and the “RAC1/PAK1/p38/MMP2” pathway, which has also been implicated in decidual inflammation, senescence and parturition (Menon *et al*., 2016). Enriched human phenotype ontology terms were related to complications of pregnancy, including “Premature Rupture of Membranes”, “Premature Birth”, “Toxemia of Pregnancy” (preeclampsia) and “Abnormal Delivery”. We also observed that several genes in the NFκB pathway, such as MYD88, CHUK, IκBKE, NFκBIE and RTKN2 were differentially expressed; NFκB signaling has been associated with the molecular etiology of preterm birth (Allport, 2001; Lindstrom and Bennett, 2005).

### HAND2 regulates IL15 expression in endometrial stromal fibroblast lineage cells

Among the genes dysregulated by *HAND2* knockdown in ESFs was *IL15* (Log_2_FC=7.98, P=7.91×10^−8^, FDR=1.49×10^−5^), a pleiotropic cytokine previously shown to be expressed in the endometrium and decidua (**Figure 3A** and **Table 1**) (Kitaya *et al*., 2000; Okada, 2000; Dunn, Critchley and Kelly, 2002; Okada *et al*., 2004; Godbole and Modi, 2010). *IL15* was robustly expressed at the first trimester maternal-fetal interface in stromal fibroblast lineage cells (**Figure 4A**) and there was a general correlation between *HAND2* and *IL15* expression in single cells (**Figure 4A inset**) (Vento-Tormo *et al*., 2018). IL15 protein localized to cytoplasm in ESF lineage cells in human pregnant decidua (**Figure 4B**) in Human Protein Atlas data. The *IL15* promoter loops to several distal sites in H3K27ac HiChIP data from E6E7hTERT endometrial cells including to regions bound by PR, NR2F2, GATA2, FOSL2, FOXO1, and SRC2, an intrinsic histone acetyltransferase that is a transcriptional co-factor of ligand-dependent hormone receptors (**Figure 4C**). The *IL15* promoter also loops to a putative enhancer in its first intron that contains a PGR binding site and SNPs marginally associated with a maternal effect on offspring birth weight (rs190663174, *P*=6×10^−4^) by GWAS (Warrington et al., 2019). *IL15* was significantly up-regulated by *in vitro* decidualization of human ESFs into DSCs by cAMP/progesterone treatment (Log_2_FC=2.15, P=2.58×10^−33^, FDR=1.59×10^−31^), and significantly down-regulated by siRNAs targeting PR (Log_2_FC=−1.24, P=6.23×10^−15^, FDR=1.80×10^−13^) and GATA2 (Log_2_FC=−2.08, P=4.16×10^−3^, FDR=0.14) (**Figure 4D**), but not NR2F2 (Log_2_FC=0.19, P=0.38, FDR=0.93) or FOXO1 (Log_2_FC=0.29, P=0.04, FDR=0.22) (**Figure 4D**). Although HAND2 binding data is not available for human stromal fibroblast lineage cells, several HAND2 binding motifs (≥0.85 motif match) are located within enhancers that loop to the *IL15* promoter.

**Figure 4.**
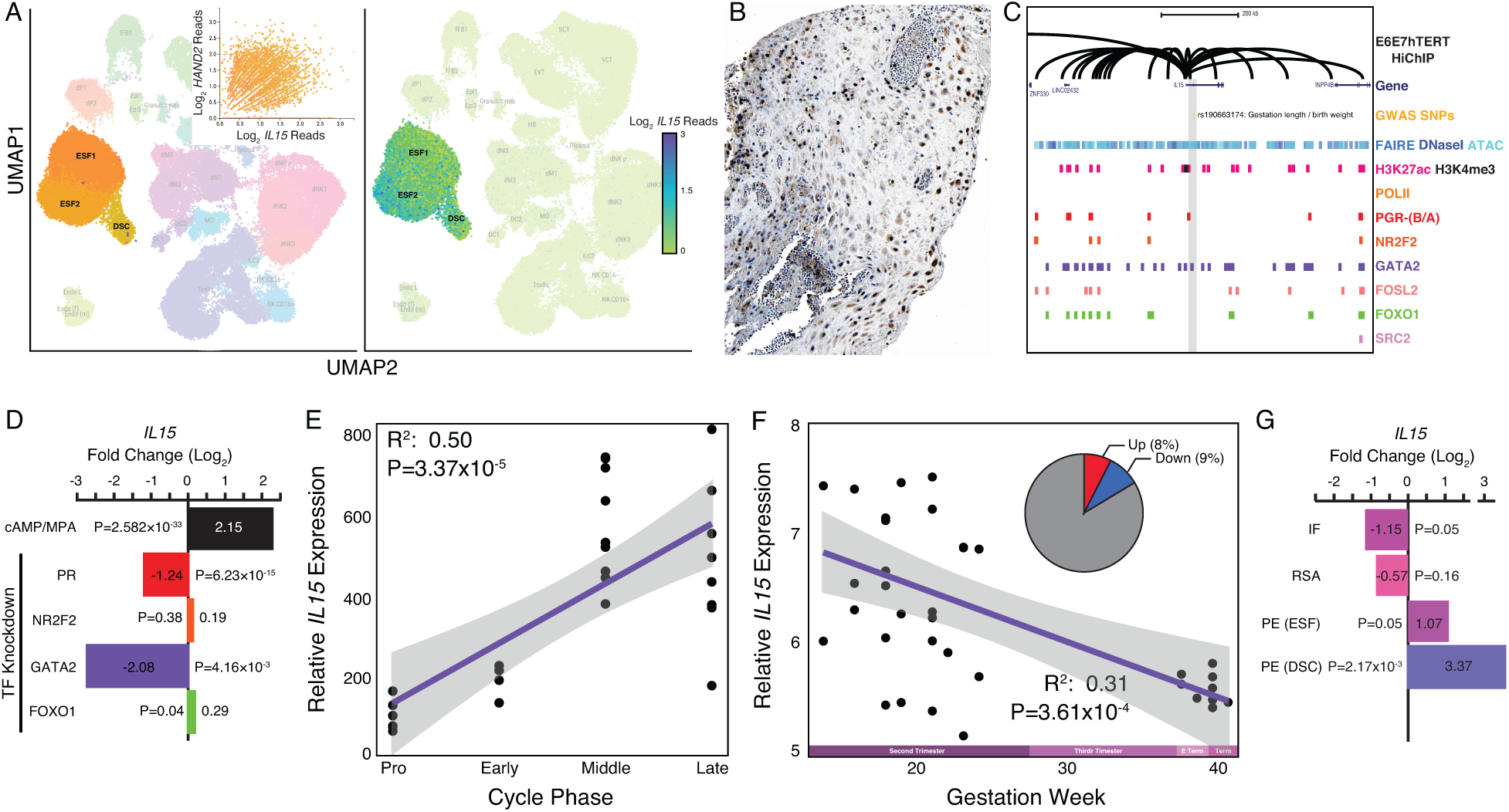
Expression of *IL15* at the maternal-fetal interface. A. UMAP clustering of scRNA-Seq data from human the first trimester maternal-fetal interface. Left, clusters colored according to inferred cell type. The ESF1, ESF2, and DSC clusters are highlighted. Inset, per cell expression of *HAND2* and *IL15* in ESF1s, ESF2s, and DSCs. Right, cells within clusters are colored according to *IL15* expression level. B. IL15 protein expression in human pregnant decidua, with strong cytoplasmic staining in endometrial stromal cells (image color auto adjusted). Image credit: Human Protein Atlas. C. Regulatory landscape of the *IL15* locus. Chromatin loops inferred from H3K27ac HiChIP, regions of open chromatin inferred from FAIRE-, DNaseI, and ATAC-Seq, and the locations of histone modifications and transcription factor ChIP-Seq peaks are shown. The location of a SNP associated with gestation length / birth weight is also shown (highlighted in gray). Note that the *IL15* promoter forms many long-range loops to regions marked by H3K27ac and bound by PR, NR2F2, GATA2, FOSL2, FOXO1, and SRC2. D. *IL15* expression is up-regulated by in vitro decidualization of ESFs into DSC by cAMP/progesterone treatment, and down-regulated by siRNA-mediated knockdown of PR and GATA2 but not NR2F2 or FOXO1. n=3 per transcription factor knockdown. H. Relative expression of *Il15* in the proliferative (n=6), early (n=4), middle (n=9), and late (n=9) secretory phases of the menstrual cycle. Note that outliers are excluded from the figure. Note that outliers are excluded from the figure but not the regression; 95% CI is shown in gray. E. Relative expression of *IL15* in the basal plate from mid-gestation to term (14-40 weeks); 95% CI is shown in gray. Inset, percent of up-(Up) and down-regulated (Down) genes between weeks 14-19 and 37-40 of pregnancy (FDR≤0.10). F. *IL15* expression is significantly up-regulated in DSCs from women with preeclampsia (PE, n=5) compared to healthy controls (n=5), while it is only marginally up-regulated in ESFs from the same patient group. It is also marginally down-regulated in the endometria of women with implantation failure (IF, n=5) and it is not differentially expressed in the endometria of women with recurrent spontaneous abortion (RSA, n=5) compared to fertile controls (n=5).

### Differential IL15 expression throughout the menstrual cycle and pregnancy

Our observations that *HAND2* is progesterone responsive and differentially expressed throughout the menstrual cycle and pregnancy suggests that *IL15*, which is controlled by HAND2 as well as cAMP/progesterone/PR/GATA2, may be similarly regulated. Indeed, like *HAND2*, we found that *IL15* expression increased as the menstrual cycle progressed, peaking in the middle-late secretory phases (**Figure 4E**) and decreased throughout pregnancy reaching a low at term (**Figure 4F**). *IL15* expression was also dysregulated in the endometria of women with infertility but not recurrent spontaneous abortion, compared to fertile controls (**Figure 4G**). In women with preeclampsia, *IL15* was not dysregulated in ESFs, but it was expressed significantly higher in DSCs, compared to controls. Thus, like *HAND2, IL15* is differentially expressed throughout the menstrual cycle and pregnancy, and in the endometria of women with infertility.

### ESF-derived IL15 promotes NK and trophoblast migration

Endometrial stromal cells promote the migration of uterine natural killer (uNK) (Chen *et al*., 2011) and trophoblast cells (Graham and Lala, 1991; Paiva *et al*., 2009; Zhu *et al*., 2009; Godbole *et al*., 2011). IL15, in particular, stimulates the migration of uNK cells (Allavena *et al*., 1997; Verma *et al*., 2000; Ashkar *et al*., 2003; Barber and Pollard, 2003; Kitaya, Yamaguchi and Honjo, 2005) and the human choriocarcinoma cell line, JEG-3 (Zygmunt *et al*., 1998). Therefore, we tested whether ESF-derived IL15 influenced the migration of primary human NK cells and immortalized first trimester extravillous trophoblasts (HTR-8/SVneo) in trans-well migration assays (**Figure 5A**). We found that ESF media supplemented with recombinant human IL15 (rhIL15) was sufficient to stimulate the migration of NK and HTR-8/SVneo cells to the lower chamber of trans-wells, compared to non-supplemented media (**Figure 5B**,**C; Supplementary Tables 9 and 10**). Conditioned media from ESFs with siRNA-mediated *HAND2* knockdown increased migration of both NK and HTR-8/SVneo compared to negative control (i.e., non-targeting siRNA) (**Figure 5B**,**C**). Conditioned media from ESFs with siRNA-mediated *IL15* knockdown reduced migration of both NK and HTR-8/SVneo cells compared to negative control (**Figure 5B**,**C**). Similarly, ESF conditioned media supplemented with anti-IL15 antibody reduced cell migration compared to media supplemented with control IgG antibody (**Figure 5B**,**C**).

**Figure 5.**
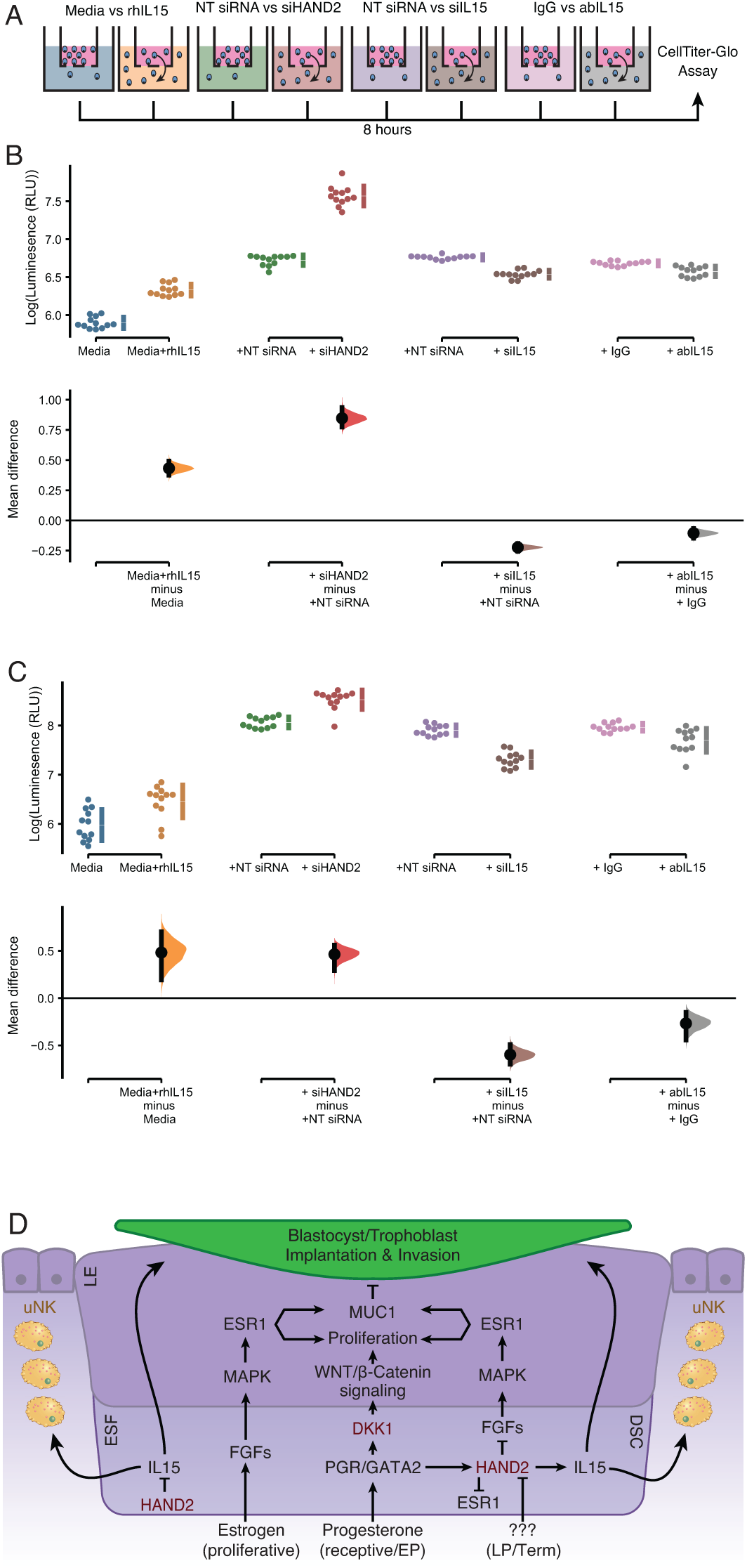
ESF-derived IL15 promotes NK and trophoblast migration in trans-well assays. A. Cartoon of trans-well migration assay comparisons. Cells that migrated to the lower chamber were quantified using the CellTiter-Glo luminescent cell viability assay after 8 hours. B. Primary natural killer (NK) cells. Raw luminescence data (RLU) is shown in the upper panel, mean difference (effect size) in experiment minus control luminescence values are shown as a dots with the 95% confidence interval indicated by vertical bars in the lower panel; distribution estimated from 5000 bootstrap replicates. The mean difference between Media and media supplemented with recombinant human IL15 (Media+rhIL15) is 0.432 [95.0% CI: 0.376 – 0.492]; P=0.00. The mean difference between ESFs transiently transfected with non-targeting siRNA (NT siRNA) and *HAND2*-specific siRNAs (siHAND2) is 0.847 [95.0% CI: 0.774 – 0.936]; P=0.00. The mean difference between ESFs transiently transfected with NT siRNA and IL15-specific siRNAs (siIL15) is −0.223 [95.0% CI: −0.256 – −0.192]; P=0.00. The mean difference between ESF media neutralized with either a non-specific antibody (IgG) or IL15-specific antibody (abIL15) is −0.106 [95.0% CI −0.147 – −0.067]; P=0.00. n=12. C. Extravillous trophoblast cell line HTR-8/SVneo. Raw luminescence data (RLU) from cells in the lower chamber is shown in the upper panel, mean difference (effect size) in experiment minus control luminescence values are shown as a dots with the 95% confidence interval indicated by vertical bars in the lower panel; distribution estimated from 5000 bootstrap replicates. The mean difference between Media and Media+rhIL15 is 0.482 [95.0% CI: 0.193 –0.701]; P=0.002. The mean difference between ESFs transiently transfected with NT siRNA and siHAND2 is 0.463 [95.0% CI: 0.291 – 0.559]; P=0.00. The mean difference between ESFs transiently transfected with NT siRNA and siIL15 is −0.598 [95.0% CI: −0.698 – 0.490]; P=0.00. The mean difference between ESF media neutralized with IgG or abIL15 is −0.267 [95.0% CI: −0.442 – −0.151]; P=0.0004. n=12. D. Model of HAND2 functions in the endometrium. During the proliferative phase HAND2 inhibits *IL15*, and thus the migration of uNK and trophoblast cells. In the receptive phase, HAND2 activates *IL15*, which promotes migration of uNK and trophoblast cells. In the receptive phase and early pregnancy (EP), HAND2 suppresses estrogen signaling by down-regulating *FGF*s and directly binding and inhibiting the ligand-dependent transcriptional activation function of ESR1. During late pregnancy/term (LP/Term), reduced HAND2 expression mitigates its anti-estrogenic functions. Parturition signal unknown (???).

## Discussion

Eutherian mammals evolved a suite of traits that support pregnancy, including an interrupted estrous cycle allowing for prolonged gestation lengths, maternal-fetal communication, implantation, maternal immunotolerance and recognition of pregnancy, and thus are uniquely afflicted by disorders of these processes. When searching for clues as to how variation in normal physiological functions can lead to dysfunction and disease, deeper understanding of the evolutionary and developmental histories of organ and tissues has the potential to provide novel insights. Here, we used evolutionary transcriptomics to identify genes that evolved to be expressed on the maternal side (endometrium) of the maternal-fetal interface during the origins of pregnancy in Eutherians, and hence may also contribute to pregnancy complications such as infertility, recurrent spontaneous abortion, and preterm birth.

Among the genes recruited into endometrial expression in Eutherians we identified *HAND2*, a pleiotropic transcription factor that plays an essential role in suppressing estrogen signaling at the time of uterine receptivity to blastocyst embedding, through its down-regulation of pro-estrogenic genes and by directly inhibiting the transcriptional activities of the estrogen receptor (Huyen and Bany, 2011; Li *et al*., 2011; Shindoh *et al*., 2014; Fukuda *et al*., 2015; Mestre-Citrinovitz *et al*., 2015; Murata *et al*., 2019). Consistent with these functions, we found evidence of estrogen activity in the endometrium of pregnant opossum. Earlier research detected similar activity in the gravid oviduct of birds and reptiles (Means *et al*., 1975; Kato *et al*., 1992; Girling, 2002; González-Morán, 2015). These data indicate that suppression of estrogen signaling during the window of uterine receptivity to implantation is an evolutionary innovation of Eutherian mammals, which involved the recruitment of *HAND2* and its anti-estrogenic functions into endometrial expression.

The roles of *HAND2* in DSCs and implantation are well-understood (Huyen and Bany, 2011; Li *et al*., 2011; Shindoh *et al*., 2014; Fukuda *et al*., 2015; Mestre-Citrinovitz *et al*., 2015; Murata *et al*., 2019; Šućurović *et al*., 2020). In contrast, the function(s) of *HAND2* at other stages of pregnancy and in ESFs remain relatively unexplored. We knocked down *HAND2* in ESFs and found that downstream dysregulated genes were enriched for human phenotype ontologies related to disorders of pregnancy, including “Premature Rupture of Membranes”, “Premature Birth”, and “Abnormal Delivery”, suggesting that *HAND2* has functions throughout pregnancy and in parturition. Indeed, we discovered that SNPs recently implicated in the regulation of gestation length and birth weight by GWAS (Warrington *et al*., 2019; Sakabe *et al*., 2020) make long-range interactions to the *HAND2* promoter. Also of note, *HAND2* expression is significantly higher in placental villous samples from idiopathic spontaneous preterm birth (isPTB) compared to term controls (Brockway *et al*., 2019). However, this difference may be related to gestational age rather than the etiology of PTB (Eidem *et al*., 2016).

Additionally, analyzing previously published datasets we noticed that *HAND2* expression decreases throughout gestation. Unlike the majority of Eutherians, where parturition closely follows the significant drop in progesterone concentrations in maternal peripheral blood, this is not the case for humans and other Old-World primates (Ratajczak, Fay and Muglia, 2010), where progesterone levels keep rising throughout gestation, reaching maximum at birth. These observations, combined with the central role of *HAND2* in mediating the anti-estrogenic actions of progesterone, suggest decreased *HAND2* at the end of pregnancy may contribute to the estrogen dominant uterine environment at the onset of labor (Pinto *et al*., 1966; Pepe and Albrecht, 1995; Mesiano *et al*., 2002; R. Smith *et al*., 2009; Ratajczak, Fay and Muglia, 2010; Welsh *et al*., 2012), despite high systemic progesterone. Low *HAND2* at the end of pregnancy in humans is therefore most likely not directly related to progesterone concentrations, suggesting that an unidentified inhibitory signal reduces endometrial *HAND2* expression. While taken collectively these data indicate a role for *HAND2* in pre/term birth, a direct mechanistic link between *HAND2* expression and the timing of parturition remains to be demonstrated.

One of the genes dysregulated by *HAND2* knockdown was the multifunctional cytokine *IL15*, which plays important roles in innate and adaptive immunity. In the context of pregnancy, it is important for the recruitment of uterine natural killer (uNK) cells to the endometrium (Kitaya *et al*., 2000; Verma *et al*., 2000; Ashkar *et al*., 2003; Barber and Pollard, 2003; Kitaya, Yamaguchi and Honjo, 2005; Laskarin *et al*., 2006). The roles of uNK cells in the remodeling of uterine spiral arteries and regulating trophoblast invasion are well known (Zygmunt *et al*., 1998; Hanna *et al*., 2006; S. D. Smith *et al*., 2009; Burke *et al*., 2010; Hazan *et al*., 2010; Lash, Robson and Bulmer, 2010; Bany, Scott and Eckstrum, 2012; Robson *et al*., 2012; Zhang, Dunk and Lye, 2013; Lima *et al*., 2014; Fraser *et al*., 2015; Felker and Croy, 2016; Renaud *et al*., 2017). Endometrial stromal cell-derived IL15 is also necessary for the selective targeting and clearance of senescent endometrial stromal cells from the implantation site by uNK cells, which is essential for endometrial rejuvenation and remodeling at embryo implantation (Brighton *et al*., 2017). Dysregulation of uNK cell-mediated clearance of these senescent cells has also been implicated in recurrent pregnancy loss (Lucas *et al*., 2020). We found that, like *HAND2, IL15* decreases throughout gestation and both genes increase in expression as the menstrual cycle progresses. Unexpectedly however, while previous studies showed that *HAND2* induces *IL15* in DSCs (Shindoh *et al*., 2014; Murata *et al*., 2020), we discovered that *HAND2* inhibited *IL15* expression in ESFs, indicating a switch in regulatory activity sometime during early menstrual cycle. These data suggest that *HAND2* regulates the appropriate timing of endometrial *IL15* expression during the menstrual cycle and pregnancy, and thus the appropriate timing of uNK cell recruitment, trophoblast migration and the clearance of senescent endometrial stromal cells from the implantation site (**Figure 5D**). Additional work is needed to elucidate the mechanisms that underlie change in *HAND2-IL15* dynamics. An intriguing possibility, however, is that progressive cell-state changes during gestation promote the transition from ESFs to DSCs to senescent DSCs (snDSCs), which have much lower *HAND2* and *IL15* expression than DSCs (Lucas *et al*., 2020). Thus, our observation of low *HAND2* and *IL15* near term may reflect a reduction of anti-inflammatory DSCs (high *HAND2* and *IL15*) and an accumulation of pro-inflammatory snDSCs (low *HAND2* and *IL15*) at the maternal-fetal interface.

Decreased *HAND2* and *IL15* expression near term and their influence on immune cells at the maternal-fetal interface may also play a role in parturition. While the signals that initiate the onset of parturition in humans and other Old-World monkeys are not known, pre/term labor is known to be associated with elevated inflammation and an influx of immune cells into utero-placental tissues (Thomson *et al*., 1999; Young *et al*., 2002; Osman *et al*., 2003; Gomez-Lopez, Guilbert and Olson, 2010; Rinaldi *et al*., 2011, 2014; Hamilton *et al*., 2012; Shynlova *et al*., 2013; Bartmann *et al*., 2014; Menon *et al*., 2016; Peters *et al*., 2016; Wilson and Mesiano, 2020). uNK cells are abundant throughout gestation (Bulmer *et al*., 1991; King *et al*., 1991; Moffett-King, 2002; Williams *et al*., 2009; Bartmann *et al*., 2014), but whether they play a role in late pregnancy and parturition is unclear. However, depletion of uNK cells rescues LPS-induced preterm birth in *IL10*-null mice (Murphy *et al*., 2005), indicating they contribute to infection/inflammation-induced preterm parturition (Murphy *et al*., 2009). CD16^+^CD56^dim^ (cytotoxic) uNK cells have also been observed in the decidua and the placental villi of women with preterm but not term labor, suggesting an association between dysregulation of uNK cells and preterm birth in humans (Gomaa *et al*., 2017). uNK cells are associated with other pregnancy complications in humans such as fetal growth restriction, preeclampsia, and recurrent spontaneous abortion (Moffett, Regan and Braude, 2004; Hiby *et al*., 2010; Wallace *et al*., 2013, 2014; Kieckbusch *et al*., 2014). Taken together, these data indicate that uNK cells may act downstream of *HAND2-IL15* signaling in the timing of parturition.

### Conclusions

Here we show that *HAND2* evolved to be expressed in endometrial stromal cells in the Eutherian stem-lineage, coincident with the evolution of suppressed estrogen signaling during the window of implantation and an interrupted reproductive cycle during pregnancy, which necessitated a means to regulate the length of gestation. Our data suggest that *HAND2* may contribute to the regulation of gestation length by promoting an estrogen dominant uterine environment near term and through its effect on IL15 signaling and uNK cell function. To further expand our understanding of *HAND2* functions at the molecular mechanistic level, multiple technical and ethical difficulties associated with studying human pregnancy *in vivo* will need to be overcome. Therefore, recently developed organoid models of the human maternal-fetal interface (Rinehart, Lyn-Cook and Kaufman, 1988; Boretto *et al*., 2017; Turco *et al*., 2017, 2018; Marinić and Lynch, 2019), which allow for *in vitro* 3D manipulation, will prove instrumental.

## Methods

### Endometrial gene expression profiling and ancestral transcriptome reconstruction

#### Data Collection

We obtained previously generated RNA-Seq data from pregnant endometria of amniotes by searching NCBI BioSample, Short Read Archive (SRA), and Gene Expression Omnibus (GEO) databases for anatomical terms referring to the portion of the female reproductive tract (FRT), including “uterus”, “endometrium”, “decidua”, “oviduct”, and “shell gland”, followed by manual curation to identify those datasets that included the FRT region specialized for maternal-fetal interaction or shell formation. Datasets that did not indicate whether samples were from pregnant or gravid females were excluded, as were those composed of multiple tissue types. Species included in this study and their associated RNA-Seq accession numbers are included in **Supplementary Table 1**.

#### New RNA-Seq data

Endometrial tissue samples from the pregnant uteri of baboon (x3), mouse (x3), hamster (x3), bat (x2), and squirrel (x2) were dissected and mailed to the University of Chicago in RNA-Later. These samples were further dissected to remove myometrium, luminal epithelium, and extra-embryonic tissues, and then washed 3x in ice cold PBS to remove unattached cell debris and red blood cells. Total RNA was extracted from the remaining tissue using the RNeasy Plus Mini Kit (74134, QIAGEN) per manufacturer’s instructions. RNA concentrations were determined by Nanodrop 2000 (Thermo Scientific). A total amount of 2.5μg of total RNA per sample was submitted to the University of Chicago Genomics Facility for Illumina Next Gen RNA sequencing. Quality was assessed with the Bioanalyzer 2100 (Agilent). A total RNA library was generated using the TruSEQ stranded mRNA with RiboZero depletion (Illumina) for each sample. The samples were fitted with one of six different adapters with a different 6-base barcode for multiplexing. Completed libraries were run on an Illumina HiSEQ2500 with v4 chemistry on 2 replicate lanes for hamster and 1 lane for everything else of an 8 lane flow cell, generating 30-50 million high quality 50bp single-end reads per sample.

#### Multispecies RNA-Seq analyses

Kallisto version 0.42.4 was used to pseudo-align the raw RNA-Seq reads to genomes (see **Supplementary Table 1** for reference genome assemblies). We used default parameters bias correction, and 100 bootstrap replicates. Kallisto outputs consist of transcript abundance estimates in transcripts per million (TPM), which were used to determine gene expression.

#### Ancestral transcriptome reconstruction

We previously showed that genes with TPM ≥ 2.0 are actively transcribed in endometrium while genes with TPM < 2.0 lack hallmarks of active transcription such as promoters marked with H3K4me3 (Wagner, Kin and Lynch, 2012, 2013). Based on these findings, we transformed values of transcript abundance estimates into discrete character states, such that genes with TPM ≥ 2.0 were coded as expressed (state=1), genes with TPM < 2.0 were coded as not expressed (state=0), and genes without data in specific species coded as missing (state=?). The binary encoded endometrial gene expression dataset generally grouped species by phylogenetic relatedness, suggesting greater signal to noise ratio than raw transcript abundance estimates. Therefore, we used the binary encoded endometrial transcriptome dataset to reconstruct ancestral gene expression states and trace the evolution of gene expression gains (0 → 1) and losses (1 → 0) in the endometria across vertebrate phylogeny (**Figure 1A**). We used Mesquite (v2.75) with parsimony optimization to reconstruct ancestral gene expression states, and identify genes that gained or lost endometrial expression. Expression was classified as an unambiguous gain if a gene was not inferred as expressed at a particular ancestral node (state 0) but inferred as expressed (state 1) in a descendent of that node, and vice versa for the classification of a loss of endometrial expression. We thus identified 149 genes that unambiguously evolved endometrial expression in the stem-lineage of Placental mammals (**Supplementary Table 2**).

#### Pathway enrichments

We used WebGestalt v. 2019 (Liao *et al*., 2019) to determine if the 149 identifies genes were enriched in ontology terms using over-representation analysis (ORA). A key advantage of WebGestalt is that it allowed for the inclusion of a custom background gene list, which was the set of 21,750 genes for which we could reconstruct ancestral states, rather than all annotated protein-coding genes in the human genome. We used ORA to identify enriched terms for three pathway databases (KEGG, Reactome, and Wikipathway), the Human Phenotype Ontology database, and a custom database of genes implicated in preterm birth by GWAS. The preterm birth gene set was assembled from the NHGRI-EBI Catalog of published genome-wide association studies (GWAS Catalog), including genes implicated in GWAS with either the ontology terms “Preterm Birth” (EFO_0003917) or “Spontaneous Preterm Birth” (EFO_0006917), as well as two recent preterm birth GWAS (Warrington *et al*., 2019; Sakabe *et al*., 2020).

### Immunohistochemistry (IHC)

Endometrial tissue from pregnant opossum (12.5 d.p.c.) was fixed in 10% neutral-buffered formalin, paraffin-embedded, sectioned at 4μm, and mounted on slides. Paraffin sections were dried at room temperature overnight and then baked for 12h at 50°C. Prior to immunostaining, de-paraffinization and hydration were done in xylene and graded ethanol to distilled water. During hydration, a 5mins blocking for endogenous peroxidase was done in 0.3% H_2_O_2_ in 95% ethanol. Antigen retrieval was performed in retrieval buffer pH6, using a pressure boiler microwave as a heat source with power set to full, allowing retrieval buffer to boil for 20mins, and then cooled in a cold water bath for 10mins. To stain sections, we used the Pierce Peroxidase IHC Detection Kit (Cat. 36000) following the manufacturers protocol. Briefly, uterine sections were incubated at 4°C overnight with polyclonal antibodies against HAND2 (Santa Cruz SC-9409), MUC1 (Novus Biologicals NB120-15481), p-ERα (Santa Cruz SC-12915), p-Erk1/2 (also known as MAPK1/2; Santa Cruz SC-23759-R) at 1:1000 dilution in blocking buffer. The next day sections were washed 3x in wash buffer, and incubated with HRP-conjugated rabbit anti-mouse IgG (H+L) secondary antibody (Invitrogen cat # 31450) at 1:10000 dilution in blocking buffer. After 30mins at 4°C slides were washed 3x in wash buffer. Slides were developed with 1x DAB/metal concentrate and stable peroxide buffer for 5mins, then rinsed 3x for 3mins in wash buffer, and mounted with Permount (SP15-100; Thermo Fisher Scientific).

### Expression of *HAND2* and *IL15* at the maternal-fetal interface

We used previously published single-cell RNA-Seq (scRNA-Seq) data from the human first trimester maternal-fetal interface (Vento-Tormo *et al*., 2018) to determine which cell types express *HAND2* and *IL15*. The dataset consists of transcriptomes for ∼70,000 individual cells of many different cell types, including: three populations of tissue resident decidual natural killer cells (dNK1, dNK2, and dNK3), a population of proliferating natural killer cells (dNKp), type 2 and/or type 3 innate lymphoid cells (ILC2/ILC3), three populations of decidual macrophages (dM1, dM2, and dM3), two populations of dendritic cells (DC1 and DC2), granulocytes (Gran), T cells (TCells), maternal and lymphatic endothelial cells (Endo), two populations of epithelial glandular cells (Epi1 and Epi2), two populations of perivascular cells (PV1 and PV2), two endometrial stromal fibroblast populations (ESF1 and ESF2), and decidual stromal cells (DSCs), placental fibroblasts (fFB1), extravillous-(EVT), syncytio-(SCT), and villus-(VCT) cytotrophoblasts (**Figure 2A**). Data were not reanalyzed, rather previously analyzed data were accessed using the cell×gene website available at https://maternal-fetal-interface.cellgeni.sanger.ac.uk.

### Expression of HAND2 and IL15 in human decidual cells

We used previously published IHC data for HAND2 and IL15 generated from pregnant human decidua as part of the Human Protein Atlas project (http://www.proteinatlas.org/; (Uhlén *et al*., 2015)). Image/gene/data available from IL15 (https://www.proteinatlas.org/ENSG00000164136-IL15/tissue) and HAND2 (https://www.proteinatlas.org/ENSG00000164107-HAND2/tissue).

### Functional genomic analyses of the *HAND2* and *IL15* loci

#### Gene expression data

We used previously published RNA-Seq and microarray gene expression data generated from human ESFs and DSCs that were downloaded from National Center for Biotechnology Information (NCBI) Short Read Archive (SRA) and processed remotely using Galaxy platform (https://usegalaxy.org/; Version 20.01) (Afgan *et al*., 2018) for RNA-Seq data and GEO2R. RNA-Seq datasets were transferred from SRA to Galaxy using the Download and Extract Reads in FASTA/Q format from NCBI SRA tool (version 2.10.4+galaxy1). We used HISAT2 (version 2.1.0+galaxy5) (Kim, Langmead and Salzberg, 2015) to align reads to the Human hg38 reference genome using single-or paired-end options depending on the dataset and unstranded reads, and report alignments tailored for transcript assemblers including StringTie. Transcripts were assembled and quantified using StringTie (v1.3.6) (Pertea *et al*., 2015, 2016), with reference file to guide assembly and the “reference transcripts only” option, and output count files for differential expression with DESeq2/edgeR/limma-voom. Differentially expressed genes were identified using DESeq2 (version 2.11.40.6+galaxy1) (Anders and Huber, 2010; Love, Huber and Anders, 2014). The reference file for StringTie guided assembly was wgEncodeGencodeBasicV33. GEO2R performs comparisons on original submitter-supplied processed data tables using the GEOquery (Davis and Meltzer, 2007) and limma (Smyth *et al*., 2002) R packages from the Bioconductor project (https://bioconductor.org/; (Gentleman *et al*., 2004)).

Datasets included gene expression profiles of primary human ESFs treated for 48 hours with control non-targeting, PGR-targeting (GSE94036), FOXO1-targeting (GSE94036) or NR2F2 (COUP-TFII)-targeting (GSE52008) siRNA prior to decidualization stimulus for 72 hours; transfection with GATA2-targeting siRNA was followed immediately by decidualization stimulus (GSE108407). We also explored the expression of *HAND2* and *IL5* in the endometria of women with recurrent spontaneous abortion and infertility using a previously published dataset (GSE26787), as well as in the endometrium throughout the menstrual cycle (GSE4888) and basal plate throughout gestation (GSE5999); HAND2 probe 220480_at, IL15 probe 205992_s_at, *HAND2* expression from proliferative and mid-secretory endometria (GSE132713).

#### ChIP-Seq and open chromatin data

We used previously published ChIP-Seq data generated from human DSCs that were downloaded from National Center for Biotechnology Information (NCBI) Short Read Archive (SRA) and processed remotely using Galaxy (Afgan *et al*., 2018). ChIP-Seq reads were mapped to the human genome (GRCh37/hg19) using HISAT2 (Kim, Langmead and Salzberg, 2015) with default parameters and peaks called with MACS2 (Zhang *et al*., 2008; Feng *et al*., 2012) with default parameters. Samples included PLZF (GSE112362), H3K4me3 (GSE61793), H3K27ac (GSE61793), H3K4me1 (GSE57007), PGR (GSE94038), the PGR A and B isoforms (GSE62475), NR2F2 (GSE52008), FOSL2 (GSE94038), FOXO1 (GSE94037), PolII (GSE94037), GATA2 (GSE108408), SRC-2/NCOA2 (GSE123246), AHR (GSE114552), ATAC-Seq (GSE104720) and DNase1-Seq (GSE61793). FAIRE-Seq peaks were downloaded from the UCSC genome browser and not re-called.

#### Chromosome interaction data

To assess chromatin looping, we utilized a previously published H3K27Ac HiChIP dataset from a normal hTERT-immortalized endometrial cell line (E6E7hTERT) and three endometrial cancer cell lines (ARK1, Ishikawa and JHUEM-14) (O’Mara, Spurdle and Glubb, 2019). This study identified 66,092 to 449,157 *cis* HiChIP loops (5 kb–2 Mb in length) per cell line, with a majority involving interactions of over 20 kb in distance; 35%–40% had contact with a promoter and these promoter-associated loops had a median span >200 kb. Contact data were from the original publication and not re-called for this study. Regions that interacted with the *IL15* promoter were searched for HAND2 binding motifs using the JASPAR 2020 database of curated, non-redundant transcription factor binding profiles stored as position frequency matrices (PFMs) (Fornes *et al*., 2020); *IL15* promoter interacting regions were searched for the HAND2 matrix profile (matrix ID MA1638.1) (Dai and Cserjesi, 2002) using the ‘Scan” option on the JASPAR website.

### Cell culture and *HAND2* knockdown

Human hTERT-immortalized endometrial stromal fibroblasts (T-HESC; CRL-4003, ATCC) were grown in maintenance medium, consisting of Phenol Red-free DMEM (31053-028, Thermo Fisher Scientific), supplemented with 10% charcoal-stripped fetal bovine serum (CS-FBS; 12676029, Thermo Fisher Scientific), 1% L-glutamine (25030-081, Thermo Fisher Scientific), 1% sodium pyruvate (11360070, Thermo Fisher Scientific), and 1x insulin-transferrin-selenium (ITS; 41400045, Thermo Fisher Scientific). 2×10^5^ cells were plated per well of a 6-well plate and 18 hours later cells in 1750μl of Opti-MEM (31985070, Thermo Fisher Scientific) were transfected with 50nM of siRNA targeting *HAND2* (s18133; Silencer Select Pre-Designed siRNA; cat#4392420, Thermo Fisher Scientific) and 9μl of Lipofectamine RNAiMAX (133778-150, Invitrogen) in 250μl Opti-MEM. BlockIT Fluorescent Oligo (44-2926, Thermo Fisher Scientific) was used as a scrambled non-targeting RNA control. Cells were incubated in the transfection mixture for 6 hours. Then, cells were washed with PBS and incubated in the maintenance medium overnight. Cells in the control wells were checked under the microscope for fluorescence the next day. 48 hours post treatment cells were washed with PBS, trypsinized (0.05% Trypsin-EDTA; 15400-054, Thermo Fisher Scientific) and total RNA was extracted using RNeasy Plus Mini Kit (74134, QIAGEN) following the manufacturer’s protocol. The knockdown experiment was done in 3 biological replicates. To test for the efficiency of the knockdown, cDNA was synthesized from 100-200ng RNA using Maxima H Minus First Strand cDNA Synthesis Kit (K1652, Thermo Fisher Scientific) following the manufacturer’s protocol. qRT-PCR was performed using QuantiTect SYBR Green PCR (204143, QIAGEN). *HAND2* primers: fwd CACCAGCTACATCGCCTACC, rev ATTTCGTTCAGCTCCTTCTTCC. *GAPDH* housekeeping gene was used for normalization; primers fwd AATCCCATCACCATCTTCCA, rev TGGACTCCACGACGTACTCA. Samples that showed >70% KD efficiency were used for RNA-Seq.

### *HAND2* knockdown RNA-Seq analysis

RNA from knockdown and control samples were DNase treated with TURBO DNA-free Kit (AM1907, Thermo Fisher Scientific) and RNA quality and quantity were assessed on 2100 Bioanalyzer (Agilent Technologies, Inc.). RNA-Seq libraries were prepared using TruSeq Stranded Total RNA Library Prep Kit with Ribo-Zero Human (RS-122-2201, Illumina Inc.) following manufacturer’s protocol. Library quality and quantity were checked on 2100 Bioanalyzer and the pool of libraries was sequenced on Illumina HiSEQ4000 (single-end 50bp) using manufacturer’s reagents and protocols. Quality control, Ribo-Zero library preparation and Illumina sequencing were performed at the Genomics Facility at The University of Chicago.

All sequencing data were uploaded and analyzed on the Galaxy platform (https://usegalaxy.org/; Version 20.01). Individual reads for particular samples were concatenated using the “Concatenate datasets” tool (version 1.0.0). We used HISAT2 (version 2.1.0+galaxy5) (Kim, Langmead and Salzberg, 2015) to align reads to the Human hg38 reference genome using “Single-end” option, and reporting alignments tailored for transcript assemblers including StringTie. Transcripts were assembled and quantified using StringTie (v1.3.6) (Pertea *et al*., 2015, 2016), with reference file to guide assembly and the “reference transcripts only” option, and output count files for differential expression with DESeq2/edgeR/limma-voom. Differentially expressed genes were identified using DESeq2 (version 2.11.40.6+galaxy1) (Anders and Huber, 2010; Love, Huber and Anders, 2014). The reference file for StringTie guided assembly was wgEncodeGencodeBasicV33. The volcano plot was generated using Blighe K., Rana S., and Lewis M. (2018): EnhancedVolcano: Publication-ready volcano plots with enhanced colouring and labeling (available at https://github.com/kevinblighe/EnhancedVolcano).

### Trans-well migration assay

#### Choice of cells and culture

Human hTERT-immortalized ESFs (T-HESC) were selected as a model ESF cell line because they are proliferative, maintain hormone responsiveness and gene expression patterns characteristic of primary ESFs, and have been relatively well characterized (Krikun *et al*., 2004). ESFs were cultured in the maintenance medium as described above in T75 flasks until ∼80% confluent. Cryopreserved primary adult human CD56+ NK cells purified by immunomagnetic bead separation were obtained from ATCC (PCS-800-019) and cultured in RPMI-1640 containing 10% FBS and IL2 in T75 flasks for two days prior to trans-well migration assays. We used the immortalized first trimester extravillous trophoblast cell line HTR-8/SVneo (Graham *et al*., 1993), because it maintains characteristics of extravillous trophoblasts and has previously been shown to be a good model of trophoblast migration and invasion (Iacob *et al*., 2008; Paiva *et al*., 2009; Hannan *et al*., 2010). HTR-8/SVneo cells were obtained from ATCC (CRL-3271) and cultured in RPMI-1640 containing 5% FBS in T75 flasks for two days prior to trans-well migration assays.

3×10^4^ ESF cells were plated per well of a 24-well plate and 18 hours later cells were transfected in Opti-MEM with 10nM (per well) of siRNA targeting *HAND2* (s18133; Silencer Select Pre-Designed siRNA, cat#4392420; Thermo Fisher Scientific) or *IL15* (s7377; Silencer Select Pre-Designed siRNA, cat#4392420; Thermo Fisher Scientific) and 1.5μl (per well) of Lipofectamine RNAiMAX (133778-150; Invitrogen). As a negative control we used Silencer Select negative control No. 1 (4390843; Thermo Fisher Scientific). ESF cells were incubated in the transfection mixture for 6 hours. Then, ESF cells were washed with warm PBS and incubated in the maintenance medium overnight. Efficiency of the knockdown was confirmed 48 hours post-treatment by qPCR, media from each well was transferred to new 24-well plates and stored at 4°C. Total RNA from cells was extracted using RNeasy Plus Mini Kit (QIAGEN) following the manufacturer’s protocol. cDNA was synthesized from 10ng RNA using Maxima H Minus First Strand cDNA Synthesis Kit following the manufacturer’s protocol. qRT-PCR was performed using TaqMan Fast Universal PCR Master Mix 2X (4352042, Thermo Fisher Scientific), with primers for *HAND2* (Hs00232769_m1), *IL15* (Hs01003716_m1) and *Malat* (Hs00273907_s1) as a control housekeeping gene. Conditioned media from samples with >70% knockdown efficiency was used for trans-well migration assays.

Corning HTS Trans-well permeable supports were used for the trans-well migration assay (Corning, cat # CLS3398). Prior to the assays, conditioned media was warmed to room temperature and centrifuged for 3min at 1000 RPM to pellet any cells. For experiments using recombinant human IL15 (rhIL15), we added 10ng/ml (AbCam, ab259403) of rhIL15 to fresh, non-conditioned ESF media; 10ng/ml has previously been shown to induce migration of JEG-3 choriocarcinoma cells (Zygmunt *et al*., 1998). For neutralizing antibody experiments, either 1μg/ml of anti-IL15 IgG (AbCam cat # MA5-23729) or control IgG (AbCam cat # 31903) were added to non-conditioned ESF media; 1μg/ml has previously been shown to neutralize ESF-derived proteins and inhibit AC-1M88 trophoblast cell migration in trans-well assays (Gellersen *et al*., 2010, 2013). Plates were incubated with shaking at 37°C for 30min prior to initiation of migration assays. During antibody incubation, NK and HTR-8/SVneo cells were collected and resuspended in fresh ESF growth media. For the trans-well migration assay, 5×10^6^ of either NK or HTR-8/SVneo cells were added to each well of the upper chamber and either treatment or control media were added to the lower chambers. Plates were incubated at 37°C with 5% CO_2_.

After 8 hours incubation, we removed the upper plate (containing remaining NK and HTR-8/SVneo cells) and discarded non-migrated cells. 50μl from each well in the lower chamber was transferred into a single well of a 96-well opaque plate. We used the CellTiter-Glo luminescent cell viability assay (G7570 Promega) to measure luminescence, which is proportional to the number of live cells per well. Data are reported as effect sizes (mean differences) between treatment and control. Confidence intervals are bias-corrected and accelerated. The P-values reported are the likelihoods of the observed effect sizes, if the null hypothesis of zero difference is true and calculated from a two-sided permutation t-test (5000 reshuffles of the control and test labels). Cumming estimation plots and estimation statistics were calculated using DABEST R package (Ho *et al*., 2019).

## Acknowledgements

The authors would like to thank the following researchers for providing pregnant endometrial samples: G.P. Wagner (Yale University) – *Monodelphis domestica*; R. R. Behringer (The University of Texas MD Anderson Cancer Center) – *Carollia perspicillata*; B.C. Paria (Vanderbilt University School of Medicine) – *Mesocricetus auratus*; A.T. Fazleabas (Michigan State University) – *Papio anubis*; D. K. Merriman (University of Wisconsin Oshkosh) – *Ictidomys tridecemlineatus*. We are also grateful to A.M. Bamberger (University Hospital Eppendorf) for providing the trans-well migration assay protocol, D. Glubb (QIMR Berghofer Medical Research Institute) for assistance in interpreting the HiChIP assay data, R. Beaumont (University of Exeter Medical School) and R.M. Freathy (University of Exeter) for assistance with interpreting maternal-fetal birth weight GWAS data. This study was supported by a grant from the March of Dimes (March of Dimes Prematurity Research Center to principal investigator VJL) and a Burroughs Welcome Fund Preterm Birth Initiative grant (1013760, to principal investigator VJL). The funders had no role in study design, data collection and analysis, decision to publish, or preparation of the manuscript. VJL thanks the Department of Human Genetics at The University of Chicago for support during the planning and preliminary data generation phase of this work. MM thanks Michael Sulak for the help with editing the manuscript.

